# Evolutionary characteristics of intergenic transcribed regions indicate widespread noisy transcription in the Poaceae

**DOI:** 10.1101/440933

**Authors:** John P. Lloyd, Megan J. Bowman, Christina B. Azodi, Rosalie P. Sowers, Gaurav D. Moghe, Kevin L. Childs, Shin-Han Shiu

## Abstract

Extensive transcriptional activity occurring in unannotated, intergenic regions of genomes has raised the question whether intergenic transcription represents the activity of novel genes or noisy expression. To address this, we evaluated cross-species and post-duplication sequence and expression conservation of intergenic transcribed regions (ITRs) in four Poaceae species. Most ITR sequences are species-specific. Those found across species tend to be more divergent in expression and have more recent duplicates compared to annotated genes. To assess if ITRs are functional (under selection), machine learning models were established in *Oryza sativa* (rice) that could distinguish between benchmark functional (phenotype genes) and nonfunctional (pseudogenes) sequences with high accuracy based on 44 evolutionary and biochemical features. Based on the prediction models, 584 rice ITRs (8%) are classified as likely functional that tend to have conserved expression and ancient retained duplicates. However, most ITRs do not exhibit sequence or expression conservation across species or following duplication, consistent with computational predictions that suggest 61% ITRs are not under selection. We outline key evolutionary characteristics that are tightly associated with likely-functional ITRs and provide a framework to identify novel genes to improve genome annotation and move toward connecting genotype to phenotype in crop and model systems.

## INTRODUCTION

Transcriptome sequencing has led to the discovery of pervasive transcription in unannotated, intergenic space in eukaryotes, including metazoan (Bertone et al. 2004; ENCODE Project Consortium 2012; Brown et al. 2014; Boeck et al. 2016), fungal (Nagalakshmi et al. 2008), and plant species (Yamada et al. 2003; Nobuta et al. 2007; Moghe et al. 2013; Krishnakumar et al. 2015; Liu et al. 2017). While intergenic transcription may be associated with nearby genes (van Bakel et al. 2010), intergenic transcripts may also represent the activity of novel genes. These intergenic transcripts may play roles as competitive endogenous RNAs (Tan et al. 2015), *cis*-acting regulatory transcripts (Guil and Esteller 2012), and/or small protein-coding regions that can be missed by gene finding programs (Hanada et al. 2013). In addition to these possible functions, intergenic transcribed regions (ITRs) may represent the products of noisy transcription, resulting from imperfect regulation of the gene expression machinery (Struhl 2007; Moghe et al. 2013) Although some of these ITRs may ultimately provide the raw materials from which novel genes may evolve *de novo* (Carvunis et al. 2012), they are not expected to be under selection. Thus, identifying the sequence features that distinguish between functional intergenic transcripts and those that are the products of noisy transcription represents a critical task in genome biology.

Here, we adopt the “selected effect” definition of function, which stipulates that a measurable activity, e.g. transcription, is considered functional if such activity is under natural selection (Amundson and Lauder 1994; Graur et al. 2013; Doolittle et al. 2014). This definition stands in contrast to the “causal role” definition that considers any reproducible biochemical activity, such as intergenic transcription, to be functional (Amundson and Lauder 1994; ENCODE Project Consortium 2012). However, pseudogenes that are remnants of once functional genes can be expressed (Zou et al. 2009; Pei et al. 2012), indicating that sequences that are not under selection could be considered functional based on the causal role definition due to biochemical activities. Given these considerations, the causal role definition of function is inadequate for distinguishing functional ITRs from noisy ones. To establish selected effect functionality in a sequence, significant conservation in sequences or activities provides strong evidence. However, lack of significant conservation does not necessarily indicate a lack of selective pressure, which may be due to selection that is significant but too weak to be detected based on sequence conservation or because only a small stretch of a sequence is under selection (Pang et al. 2006; Ponting 2017). Instead of relying on a single line of evidence such as sequence conservation, an approach that integrates genetic, evolutionary, and biochemical evidence has been suggested (Kellis et al. 2014). Based on this framework, predictive models were established that were highly effective at identifying sequences with significant fitness cost when mutated (Gulko et al. 2014) and distinguishing human and *Arabidopsis thaliana* protein coding and RNA genes from pseudogenes (Tsai et al. 2017; Lloyd et al., 2018). Thus, integration of evolutionary and biochemical signatures could provide valuable insight in distinguishing functional and noisy ITRs.

In this study, we investigate the extent of sequence and expression conservation as well as potential functionality of ITRs using data from four Poaceae (grass) species: *Oryza sativa* (rice), *Brachypodium distachyon*, *Sorghum bicolor* (sorghum), and *Zea mays* (maize). Phylogenetically, these four species fall into two clades where one clade consists of maize and sorghum diverged ~15 million years ago (MYA) (Skendzic et al. 2007; Liu et al. 2014), and the other contains rice and *B. distachyon* diverged ~47 MYA (Massa et al. 2011). All four species share ancient whole genome duplications (WGDs) (Paterson et al. 2004; Tang et al. 2010) while the maize lineage experienced a more recent WGD ~12 MYA (Swigoňová et al. 2004). Thus, studies of ITRs in these species are well-suited to assess not only sequence and expression conservation of ITRs but also ITR duplicate retention post duplication. In addition, using rice as an example, we generated function prediction models by integrating rice mutant phenotype, sequence conservation, transcriptome, histone modification, DNA methylation, and nucleosome occupancy data to predict functional ITRs genome-wide.

## RESULTS

### Identification and classification of Poaceae transcribed regions

To investigate the properties and potential functionality of polyadenylated intergenic transcripts, we focused on four Poaceae species (**Fig. 1A**). In each of the four species, we identified transcribed regions and classified them as parts of exons, introns, or pseudogenes (see Methods). Transcriptome datasets were from developmentally-matched leaf, seed, and reproductive tissues (Davidson et al. 2011; Davidson et al. 2012; **Table S1**). Transcribed regions that did not overlap with gene or pseudogene annotation were considered as intergenic. Intergenic transcribed regions (ITRs, **Table S2)** accounted for only 4-7% of mapped reads and 6-12% of transcribed regions (defined by continuous read mapping) in each species (**Fig. 1B**). In contrast, 92-96% of mapped reads overlapped with annotated exons and/or introns (**Fig. 1B**).

**Figure 1.**
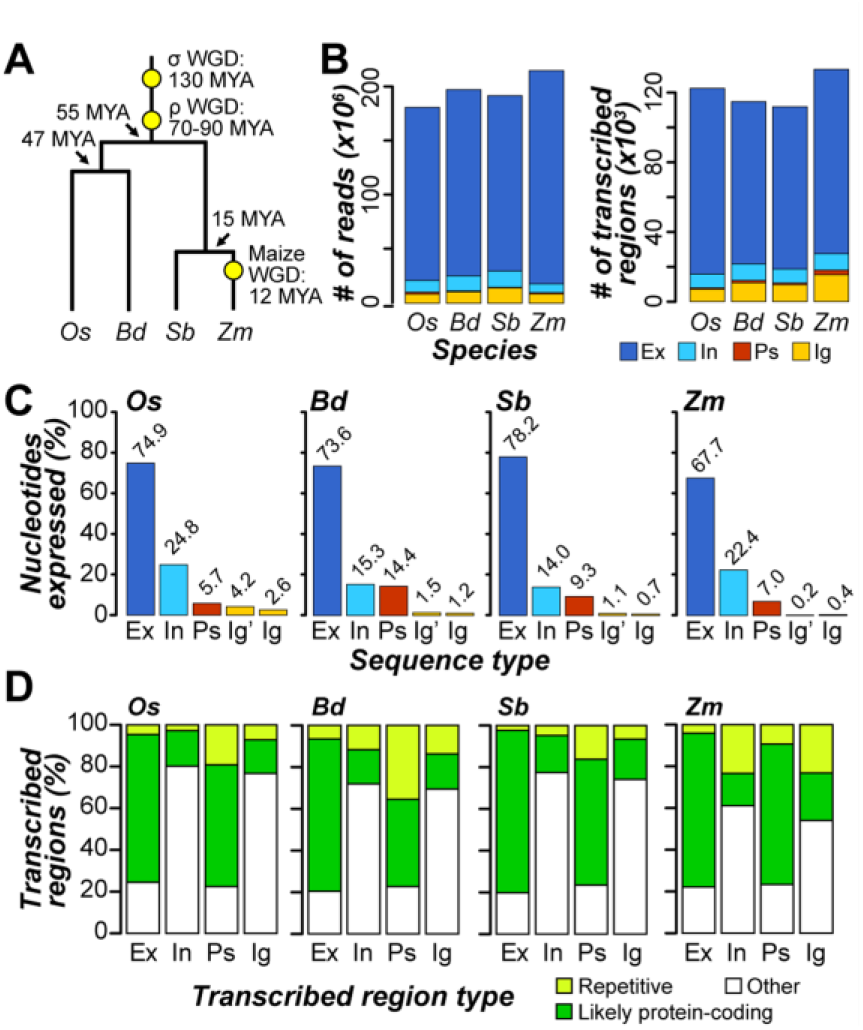
Relationships between and transcriptome content in the four Poaceae species. **(A)** Phylogenetic relationships between rice (*Os*), *B. distachyon* (*Bd*), sorghum (*Sb*), and maize (*Zm*). Whole genome duplication (WGD) events are marked with yellow circles. MYA: millions of years ago. **(B)** Number of reads mapping to (left panel) and transcribed regions overlapping (right panel) exons (Ex; dark blue), introns (In; cyan), pseudogenes (Ps; red), and intergenic regions (Ig; orange). **(C)** Percent of nucleotides overlapped with transcribed regions that are annotated as exons, introns, pseudogenes, and intergenic regions. Abbreviations are the same as in **(A)**. Ig’ represents the proportion of intergenic space covered by transcribed regions that also overlap genic or pseudogenic regions. **(D)** Percent of transcribed regions that were classified as highly repetitive (lime), likely protein-coding (green), or neither of these two (Other; white).

We also found that only 0.4-2.6% of intergenic space was covered by transcribed regions in each species (**Fig. 1C**). Thus, intergenic transcription in Poaceae is rare relative to genic expression, consistent with previous findings in *A. thaliana* (Lloyd et al. 2018). Only a small fraction of plant intergenic space is expressed compared to the ~60% intergenic expression uncovered in the human genome (ENCODE Project Consortium 2012).

Despite the relative scarcity of intergenic transcripts, we identified 7,000 to 16,000 ITRs in these four species (**Fig. 1B**, right panel, orange). We asked whether ITRs resemble known annotated sequences and found that 16-23% of ITRs in the four species have significant translated sequence similarity to plant proteins and/or contain a known protein domain (**Fig. 1D**). In addition, only 56-94 ITRs (ό1% in each species) contain known RNA gene domains. ITRs resembling annotated genes may be unannotated exons, novel genes, or pseudogenes that are still expressed. Only 7-23% of ITRs are highly repetitive (**Fig. 1D**), indicating that expression of abundant transposable element families does not account for the majority of intergenic expression in these species. Overall, 54-77% of ITRs among the four species do not resemble annotated features and are not highly repetitive.

### Expression properties of ITRs and their associations with neighboring genes

We next contrasted four expression properties – transcript fragment length, expression level and breadth, and reproducibility across experiments – between ITRs and transcribed exons. Compared to transcribed exons, ITRs in all four species are significantly shorter (Mann Whitney U tests, all *p*<7×10^-105^), expressed at lower levels (all *p*<8×10^-311^), and expressed in fewer tissues, (all *p*<2×10^-306^) (**Fig. S1A-C**). We found that 76-88% of ITRs were reproducible across leaf transcriptome biological replicates (**Fig. S1D**). Although these proportions were significantly lower than those of transcribed exons (93-95%; Fisher’s Exact Tests (FETs), all *p*<3×10^-10^), they far exceeded randomly expected percentages (2-21%; FETs, all *p*<7×10^-10^). This is indicative of the presence of hotspots in intergenic space where reproducible transcription originates, either due to the presence of truly functional sequences or noisy transcription resulting from spurious regulatory signals.

In mammalian systems, intergenic transcripts have been suggested to be associated with nearby genes (van Bakel et al. 2010), either as unannotated exon extensions or products of run-on transcription. Interestingly, the expression correlations between ITRs and proximal annotated gene neighbors (<500 bp) were significantly higher than expression correlation between ITRs and distal gene neighbors (>500bp) or between proximal gene-gene neighbors (**Fig. 2A**). There are two potential explanations for this pattern: 1) ITRs proximal to genes may be unannotated gene extensions or 2) gene regulation may influence expression patterns of nearby unrelated transcripts. Consistent with the second explanation, we found that pseudogenes, which are not gene extensions, proximal to gene neighbors exhibited similar degrees of expression correlation as proximal gene-ITR pairs (**Fig. 2**). Taken together, the expression characteristics of ITRs are consistent with noisy transcripts that are expected to be short in length, and weakly and narrowly expressed (Struhl 2007). We also find no evidence to support the notion that the majority of Poaceae ITRs represent unannotated exons of known genes. In fact, among the four Poaceae species, ITRs were significantly more distant from genes than randomly expected (U tests, all *p*<2×10^-36^; **Fig. 2B**).

**Figure 2.**
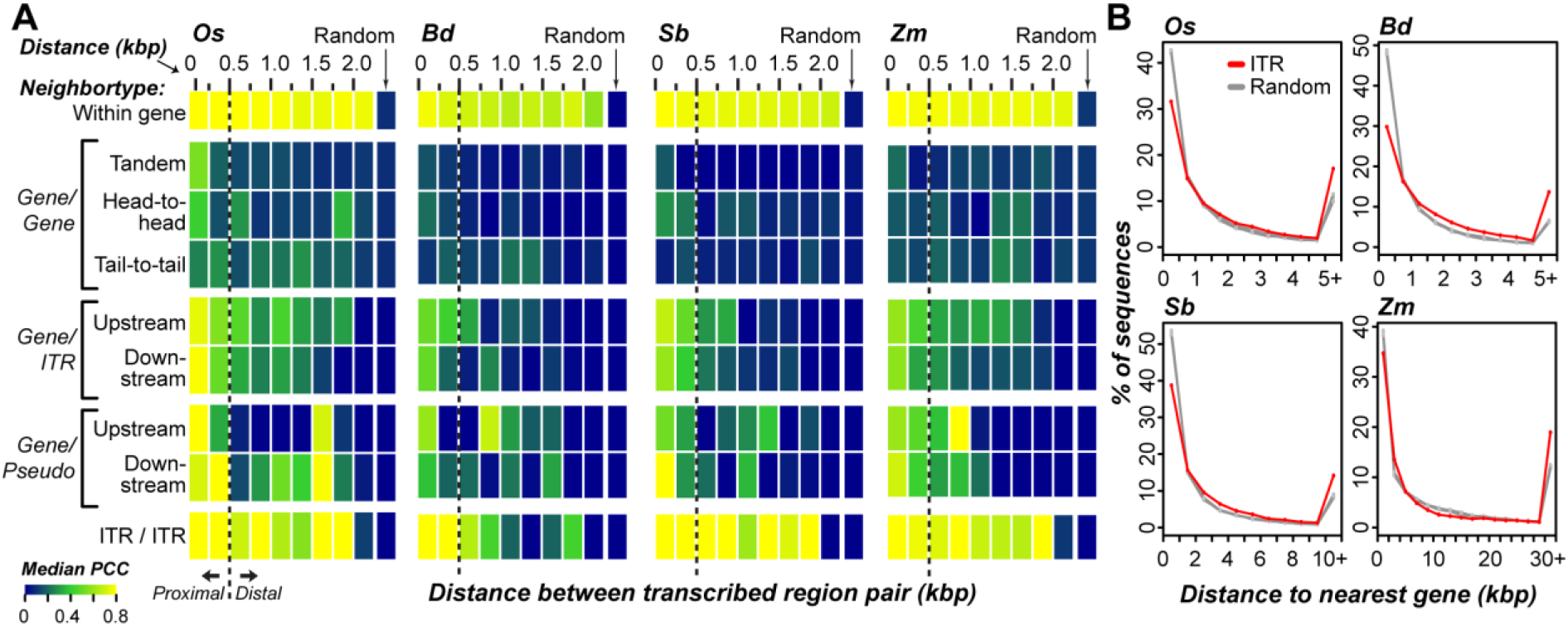
Expression correlation and distance between intergenic transcribed regions (ITRs) and neighboring genes. (**A**) Heatmaps of expression correlation between neighboring pairs of transcribed regions at various distance (kbp: kilobase pairs) in four species. Colors represent the median Pearson’s correlation coefficient (PCC) of expression levels (FPKM) across tissues between all transcribed regions pairs within a distance bin. Neighboring transcribed region pairs were classified according to whether they were in the same gene (Within gene), neighboring genes (Gene/Gene), genes and neighboring ITRs (Gene/ITR), gene and pseudogene neighbors (Gene/Pseudo), or neighboring ITRs (ITR/ITR). Random: based on 10,000 randomly-selected pairs of the same type. Neighboring gene pairs were sub-classified according to whether genes were oriented in the same direction (Tandem), or different directions with proximal 5’ (Head-to-head) or 3’ (Tail-to-tail) regions. Gene/Pseudo and Gene/ITR pairs were sub-classified according to whether the pseudogene or intergenic transcribed region was upstream or downstream of a gene neighbor. (**B**) Distance distributions between ITRs and their nearest genes (ITR) and randomly-selected intergenic regions of the same number and length distribution as ITRs in each corresponding species (five replicates, Random). Note the five randomly sampled distributions are highly overlapping.

### Cross-species sequence and expression conservation of ITRs

Sequence conservation due to selective pressure is a hallmark of functional genome regions. We considered a sequence as conserved if it exhibited cross-species sequence similarity significantly greater than that observed for random, unexpressed intergenic sequences (see Methods). Under this framework, we found that only 15-19% of ITRs among the four species were conserved across species (**Fig. 3A**). Significantly fewer ITRs were conserved across species compared to transcribed exons (79-83% conserved), introns (21-35%), or pseudogenes (32-54%; FET, all *p*<2×10^-9^). Thus, ITRs primarily represent species-specific sequences.

**Figure 3.**
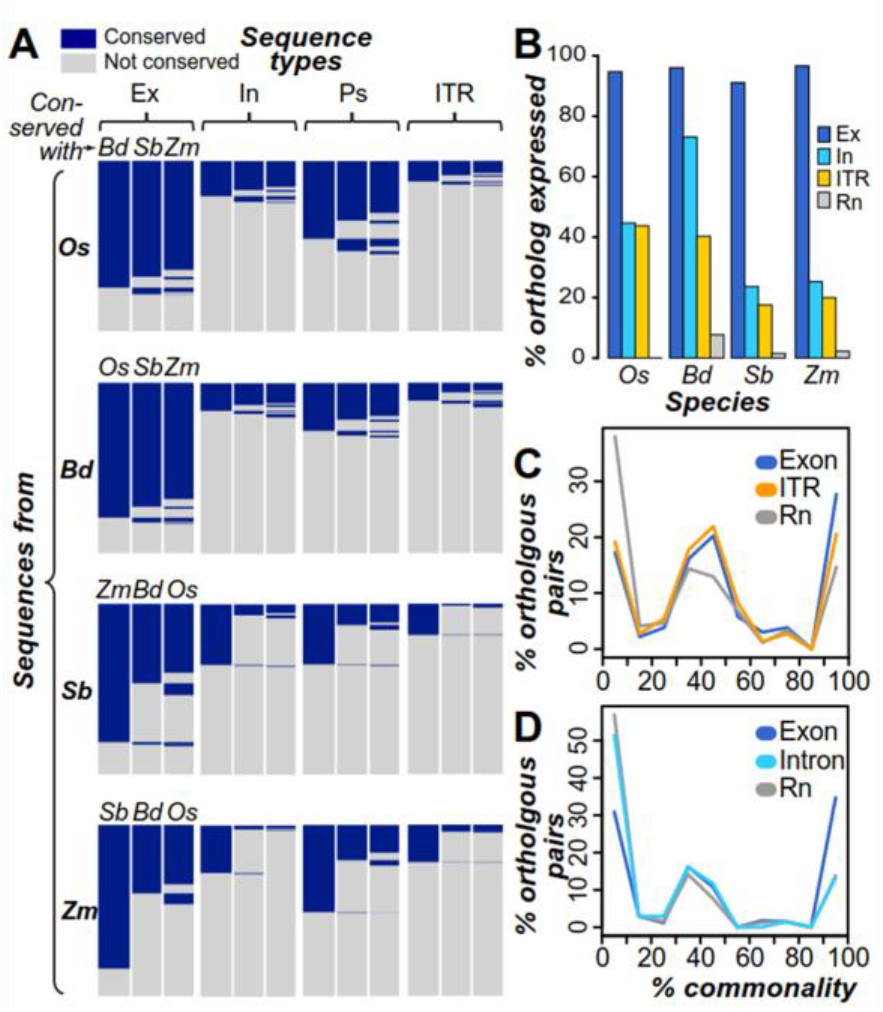
Sequence and expression conservation of transcribed regions. **(A)** Heatmaps of crossspecies sequence conservation. Blue: conserved (cross-species sequence similarity significantly greater than that observed for random, unexpressed, non-repetitive intergenic sequences). Gray: not conserved. Species abbreviations follow **Fig. 1**. Ex: exon, In: intron, Ps: pseudogene. **(B)** Percentages of syntenic orthologs with expression evidence. Syntenic orthologs were identified by conservation across pairs of cross-species syntenic blocks (see Methods). Ex: exon, In: intron **(C,D)** Percent commonality in tissue expression between pairs of cross-species homologous exons (based on similarity but not synteny), ITR (panel **C**), introns (panel **D**), and randomly-generated expression vectors (Rn). Given the influence that expression breadth exerts on expected % commonality, comparisons among orthologous transcribed exons and random expression vectors where matched in expression breadth to that observed in orthologous ITR and transcribed intron pairs (see Methods). Note that this results in distinct distributions for transcribed exons and random expression vectors for ITRs and transcribed introns.

We next evaluated the expression conservation of ITRs by two measures. First, we identified syntenic orthologs (see Methods) and determined whether regions in other species orthologous to ITRs were also expressed. We found that 18-44% of conserved ITRs and their orthologs were both expressed, significantly lower than the percentages for transcribed exons and their orthologs (91-97%, FET, all *p*<2×10^-10^; **Fig 3B**). Thus, species-specific gain of expression may explain ITR expression. Alternatively, if the ancestral ITR sequence was expressed, maintenance of ITR expression across species is rare. However, significantly fewer unexpressed intergenic sequences had an ortholog that was expressed (0-7%) compared to ITRs (FET, all *p*<0.003), indicating that, when both an ITR and its ortholog are expressed, this expression conservation likely represents non-random events that are under selection. ITRs with significant similarities to protein coding genes (**Fig. 1D**) are more likely to have conserved expression across species, as 32% of ITRs with similarity to protein-coding sequences were associated with an expressed ortholog compared to 11% of ITRs without protein similarity (FET, *p*<3×10^-6^). We also identified sequence blocks conserved across all four species (see Methods). Among 639 blocks that were intergenic in all species, we found that 15% were expressed in at least one species (**Fig. S2A**), compared to >99% of blocks composed of exons in all four species (FET, *p*<6×10^-11^; **Fig. S2B**). When an intergenic conserved sequence block was expressed, the expression was limited to one species in 74% of cases. While expression in two, three, or four species was documented in only 18%, 6%, and 1.3% (n=1) cases, respectively (**Fig. S2A**), compared to 74% of exon blocks that were expressed in all four species (**Fig. S2B**). Overall, we find that the two analyses lead to the same conclusion: the few ITRs with sequence conservation rarely exhibit conserved expression.

For the second measure of expression conservation, we evaluated whether ITRs and their expressed orthologs were expressed in similar tissues. For a pair of orthologs, we calculated percentage of tissue commonly expressed (% commonality, see Methods). Orthologous pairs of ITRs had an average of 48% commonality, which was lower than the average for orthologous transcribed exons (53%; U test, *p*=0.04), but significantly higher than the average for random sequence pairs (36%, *p*<6×10^-6^; **Fig. 3C**). By comparison, expressed tissue % commonality between transcribed introns and their orthologs (mean % commonality=27%) was no different than random (mean=26%; U test, *p*=0.33; **Fig. 3D**). These findings suggest that ITRs with cross-species expression conservation, particularly those with higher % tissue commonality, may be enriched for sequences under selection.

### Sequence and expression conservation of ITR paralogs

In the previous section, we examined cross-species sequence and expression conservation. Considering that plant genomes harbor a rich history of large- and small-scale duplication events, we next asked whether ITR duplicates tend to be retained over time, and if so, whether the paralogous ITRs have similar expression patterns. We excluded highly-repetitive sequences (**Fig. 1D**) from the analyses because they were duplicated by definition. We found that 47-50% of rice, *B. distachyon*, and sorghum ITRs were duplicated, compared to 45-48% of transcribed exons and 31-66% of random, unexpressed intergenic sequences (**Fig. S3**). In maize, 82% of ITRs, 74% of transcribed exons, and 95% of random, unexpressed intergenic sequences were duplicated (**Fig. S3**). The high percentage of duplicated sequences from random, unexpressed intergenic regions in maize is due to an overall greater number of paralogs identified among maize exons and repetitive sequences, which were used to define thresholds to call a sequence as repetitive (see Methods). Based on the estimated base substitution rates (*K*) as a proxy for time (**Fig. 4A**), ITR duplicates in all species were generated from more recent duplication events compared to transcribed exons (U tests, all *p*<6×10^-127^),instead more closely resembling introns and pseudogenes that experience weaker or no selection. Together with the rarity of ITR duplicates at larger *K* (>0.06, **Fig. 4A**) compared to exons in annotated genes, most ITR duplicates are likely not maintained.

**Fig. 4.**
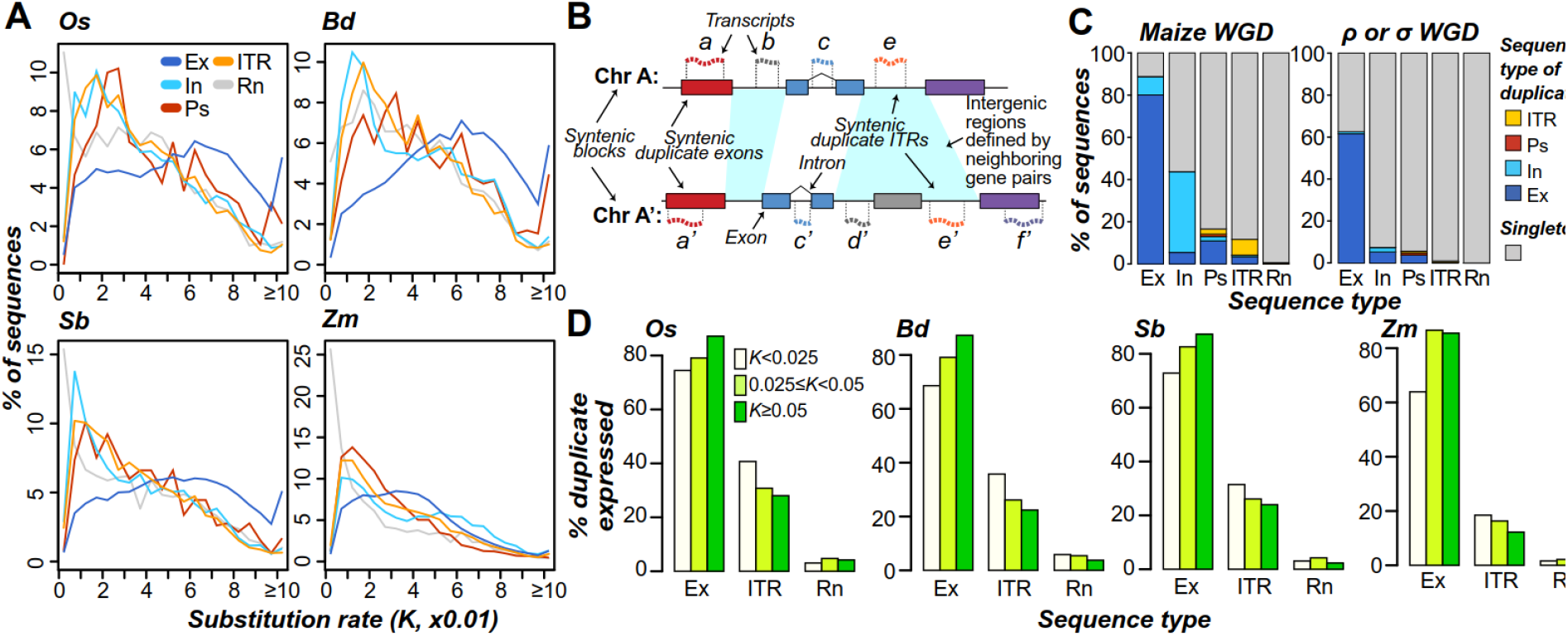
Duplication characteristics of transcribed regions. **(A)** Distributions of nucleotide substitution rates (*K*) between a sequence and its top within-species nucleotide sequence match. **(B)** Schematic of syntenic duplicate identification method. Syntenic regions from two chromosomes (Chr A and A’) are shown, with duplicate genes indicated by boxes with matching colors and transcripts by wavy, dashed lines. Cyan: intergenic region pair defined by neighboring duplicate pairs. Syntenic duplicates are defined by their expression state and sequence types, including transcribed exon pair (a and a’), transcribed intron pair (c and c’), and ITR pair (e and e’). Also shown are examples of transcribed regions that lack a syntenic duplicate: b, d’, and f’. Note that syntenic paralogs were also identified among non-transcribed sequences in the same manner. For example, if e’ was not expressed, a query with the sequence underlying e would identify similarity to the sequence underlying e’. **(C)** Proportions of syntenic duplicates from the maize lineage-specific whole genome duplication (Maize WGD; left panel) and ρ/σ WGDs (right panel). Singleton: sequences that have no syntenic duplicate identified. Ps: pseudogene. Other sequence types are abbreviated as in **Fig. 3C,D**. **(D)** Percent of expressed duplicates among sequence types in three nucleotide substitution rate (*K*) bins. Species abbreviations follow **Fig. 1**. Sequence type abbreviations follow **Fig. 3**.

To more definitively determine the relationship between the timing of duplication events and ITR duplicate retention, we identified ITRs present in duplicated genome blocks derived from whole genome duplication (WGD) events (referred as syntenic blocks, **Fig. 4B**; **Fig. S4**). We first looked at duplicate sequences derived from a WGD event 12 MYA in maize (Swigoňová et al. 2004; see Methods). Among the 4,906 non-repetitive maize ITRs in these syntenic blocks, 463 (9%) had syntenic duplicates (**Fig. 4C**), significantly more than that of random, unexpressed intergenic regions (2%; FET, *p*<4×10^-10^) but significantly fewer than protein-coding genes (36%, FET, *p* < 3×10^-16^). Thus, as many as 91% of ITR duplicates were deleted, mutated beyond recognition, or translocated within a span of 12 million years. When considering syntenic blocks from the much older ρ and σ WGD events >70 MYA in all four species, only 1.0% (49 of 4,859) of ITRs had syntenic duplicates, compared to 28-31% of protein-coding genes, indicating an overall lack of ancient duplicates. Nonetheless, a greater proportion of ITRs have ρ or σ WGD-derived duplicates compared to random, unexpressed intergenic regions (0.2%; FET, *p*<0.002; right panel, **Fig. 4C**), suggesting that a subset of these ITR duplicates have been retained for over 70 MY.

We next assessed whether expression was conserved among duplicate ITRs and found that 18-34% ITR duplicates (most similar paralogs) were also expressed depending on the species, compared to 79-84% of paralogs of transcribed exons (FET, all *p*<6×10^-10^). In addition, ITR paralogs generated from recent duplication events are more likely to be expressed than those from older events (U tests, all *p*<3×10^-5^; **Fig. 4D**). This suggests that the expression of ITRs may be initially maintained following duplication, but frequently lost over time. By contrast, older duplicates of transcribed exons are more frequently expressed than recent duplicates (U tests, all *p*<8×10^-109^; **Fig. 4D**). Compared to ITR duplicates, fewer duplicates of random unexpressed intergenic sequences were expressed (2-7%; FET, all *p*<4×10^-10^). Overall, ITRs were duplicated nearly as often as annotated genes but significantly fewer ITR paralogs were maintained over time or expressed compared to paralogs of annotated genes. In addition, the negative relationship between timing of duplication and percent expressed ITR paralogs suggest transcription of ITR duplicates tend to be lost over time, characteristics that could be indicative of sequences that are not under selection.

### Predicting the functionality of rice transcribed regions

To further evaluate the selected effect functionality of ITRs, we first examined which evolutionary and biochemical characteristics can distinguish whether a transcribed region resembled a phenotype gene or pseudogene. Due to data availability, we focused solely on rice. For benchmark functional sequences, we identified 513 transcribed regions that overlap exons of genes with documented loss-of-function phenotypes (referred to as phenotype exons, **Table S3**). Benchmark nonfunctional sequences consisted of 262 transcribed regions that overlapped pseudogenes but not annotated genes (transcribed pseudogenes, **Table S3**). To distinguish between the benchmark functional and nonfunctional sequences, 44 evolutionary and biochemical features in five categories were examined including transcription activity, sequence conservation, DNA methylation, histone modifications, and nucleosome occupancy (**Table S3, 4**; see Methods). Among them, 38 features had significantly different value distributions between phenotype exons and transcribed pseudogenes (all *p*<0.03) but the effect sizes (differences in values) tend to be small. That is, if we used any single feature to build a naïve classifier, the median Area Under Curve-Receiver Operating Characteristic (AUC-ROC) = 0.62 (**Table S4**).

Although these naïve classifiers perform better than random guessing (AUC-ROC=0.5), they were far from perfect (AUC-ROC=1). Therefore, we considered all 44 features in combination using a machine learning approach to generate a function prediction model (see Methods). The resulting model could distinguish between phenotype exons and transcribed pseudogenes with high AUC-ROC (0.94; **Fig. 5A**) and precision/recall (**Fig. 5B**). The model provides a score between 0 and 1 (referred to as functional likelihood) where higher and lower scores indicate similarity to phenotype exons and transcribed pseudogenes, respectively (**Fig. 5C-H**, **Table S5**). Using a stringent functional likelihood threshold (T1=0.60) where only 5% of pseudogenes are misclassified as functional, 75% of phenotype genes are correctly predicted as functional (**Fig. 5C**). We also defined a second threshold (T2=0.29) where 5% of phenotype exons are misclassified as pseudogenes and the majority of pseudogenes (70%) remain correctly predicted (**Fig. 5D**). These findings indicate that the functional prediction model can distinguish between functional and nonfunctional sequences. Given most of the benchmark phenotype exons were protein-coding, we applied the functional prediction model to 22 transcribed miRNAs. Most transcribed miRNAs (64%) had functional likelihood values between T1 and T2 and cannot be confidently classified as either phenotype exon-like or pseudogene-like (**Fig. 5E**), indicating that the idiosyncrasies of miRNA sequences are not completely captured by the model. However, we cannot rule out the possibility of false-positive annotations present in the set of transcribed miRNAs. Nevertheless, most miRNA sequences (73%) are not predicted as pseudogene-like (i.e. functional likelihood < T2).

**Fig. 5.**
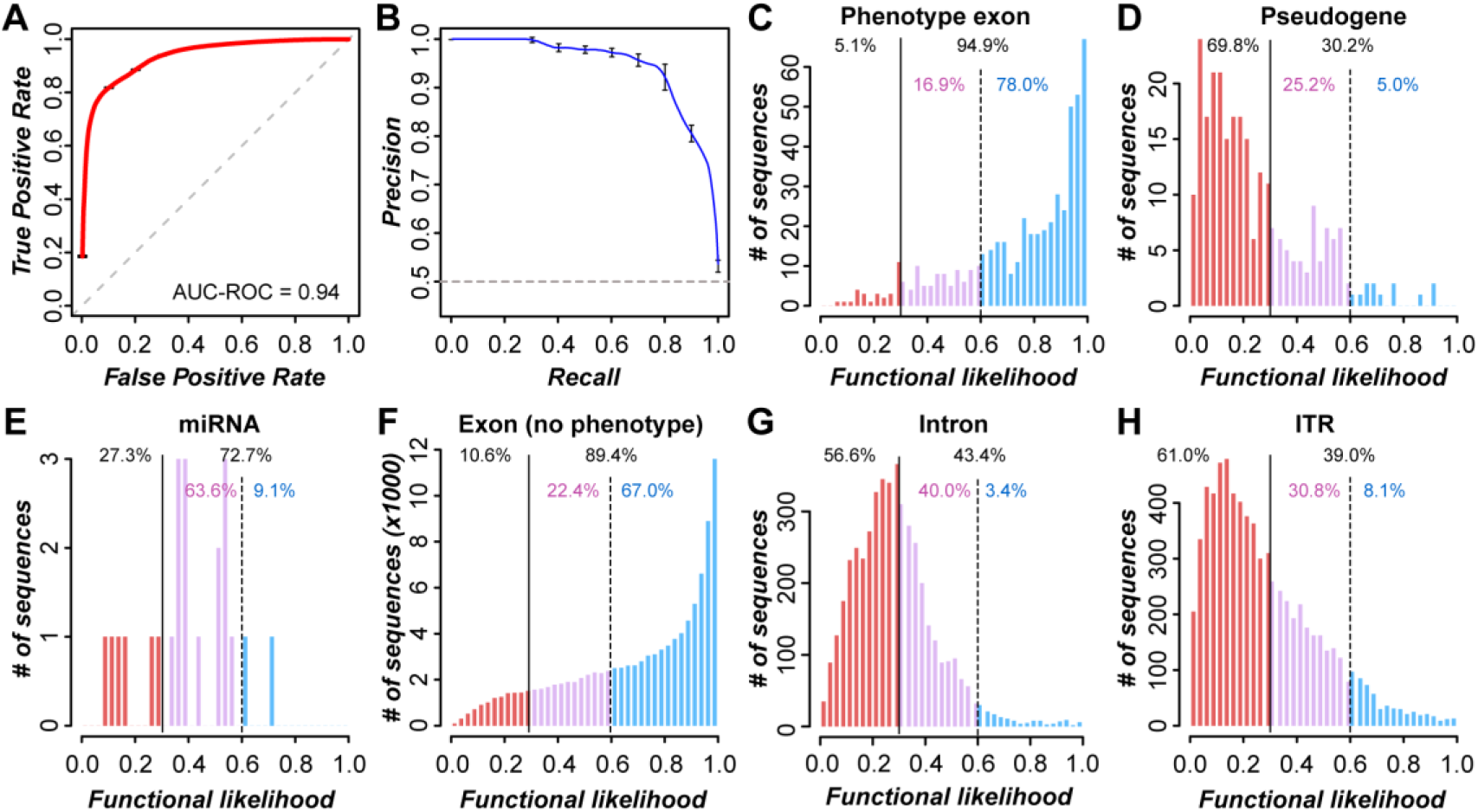
Machine learning-based functional prediction. (**A**) Receiver operating characteristic (ROC) curve for predictions of phenotype exons and transcribed pseudogenes. AUC-ROC: area under curve-ROC. (**B**) Precision-recall curve for the predictions from (**A**). Gray dashed lines **(A)** and **(B)**: random expectation. **(C-H)** Histograms of functional likelihood scores for phenotype exons (**C**), pseudogenes (**D**), transcribed miRNAs (**E**), transcribed exons without phenotype evidence (**F**), transcribed introns that are not alternatively spliced (**G**), and ITRs (**H**). Dotted black line: threshold T1 based on a 5% false positive rate. Solid black line: threshold T2 based on a 5% false negative rate. Blue, purple, and red: phenotype exon-like, ambiguous, and transcribed pseudogene-like, respectively.

We applied the function prediction model to all rice transcribed regions that were not used to establish the models, including transcribed exons, transcribed introns, and ITRs. Among transcribed exons, 67% were predicted as functional based on T1 and 11% were pseudogene-like based on T2 (**Fig. 5F**). Among transcribed introns that are not annotated as alternatively spliced exons, only 3% were predicted as functional based on T1 and the majority (57%) were predicted as pseudogene-like based on T2 (**Fig. 5G)**, indicating that the functional prediction model can effectively distinguish between exon and intron sequences. Next, we applied the prediction models to ITRs and found that 8% were predicted as functional based on T1 and 61% were pseudogene-like based on T2 (**Fig. 5H**). We found that there was a significant negative correlation between ITR functional likelihoods and their distance to neighboring genes (Spearman’s rank correlation, ρ = −0.17, *p*<3×10^-47^), with notably higher likelihoods among ITRs <500 bp from a gene (**Fig. S5**). It remains unclear whether functional predictions among ITRs are influenced by the chromatin state of neighboring genes or these ITRs represent truly functional sequences. Ultimately, the majority of rice ITRs (61%) are predicted as non-functional based on T2, which is consistent with the proportion of potentially-functional ITRs in *A. thaliana* (Lloyd et al. 2018). By and large, the functional likelihood distribution of ITRs are similar to those of pseudogenes and introns, but distinct from miRNA (peaking at middle functional likelihood) and phenotype exons. These findings suggest that ITRs may primarily result from transcriptional noise, although a small percentage are likely novel genes.

### Identification of evolutionary characteristics that closely associate with functional sequences

With subsets of ITRs predicted as functional and nonfunctional, we next sought out how well sequence and expression conservation characteristics correlated with functional predictions. Among rice ITRs with cross-species homologs in *B. distachyon*, sorghum, or maize (defined based on sequence similarity, not synteny). Among rice ITRs with cross-species homologs, 36% were predicted as functional based on T1, significantly higher than 3% for ITRs without putative homologs (Fisher’s Exact Test, *p*<2.2×10^-16^, **Fig. 6A**, left column), although lower than that for exons (80%; FET, *p*<2.2×10^-16^, **Fig.6A**, right column). We next tested whether rice ITRs with expressed orthologs (defined based on synteny) in *B. distachyon* (diverged ~47 MYA) would primarily represent functional sequences and found that 50% of ITRs with expressed orthologs were predicted as functional based on T1, significantly higher than ITRs with non-expressed orthologs (12%; FET, *p*<0.05; **Fig. 6B**, left column). Similarly, 81% of transcribed exons with an expressed ortholog were predicted as functional based on T1 (**Fig. 6B**, right column). Since cross-species homology was used as a feature for model building, the correlation is not surprising. But expression of an ortholog was not used as a feature, indicating that our findings provide strong support for ITR functionality as defined by the machine learning model. We next investigated the relationship between functional likelihood of ITRs/exons and duplication status. We found that single-copy ITRs exhibited significantly higher functional likelihood (median = 0.26) compared to ITR duplicates (median = 0.21; U test, *p*<3×10^-16^; **Fig. 6C**, left column), but lower than those of singleton transcribed exons (median = 0.78, U test, *p*<3×10^-16^, **Fig. 6C**, right column). We also found that ITRs with expressed paralogs tend to have higher functional likelihood (median= 0.25) compared to those with unexpressed paralogs (median = 0.18; U test, *p*<3×10^-16^; **Fig. 6D**). Despite significant correlation with functional likelihood, only 9% and 10% of single-copy ITRs and ITRs with an expressed paralog, respectively, were predicted as functional based on T_1_.

**Fig. 6.**
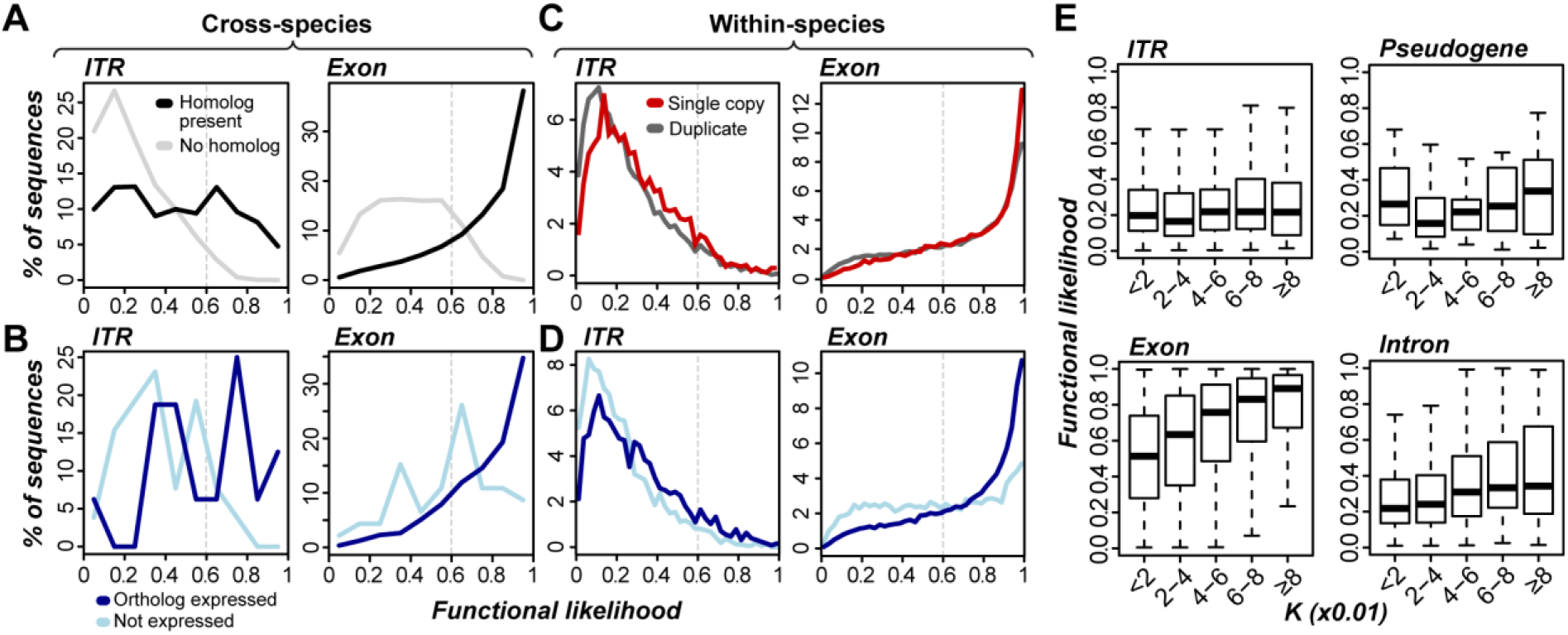
Relationships between functional likelihood scores and sequence/expression conservation characteristics. (**A**) Functional likelihood distributions for ITRs (left) and transcribed exons (right) with (black) or without (gray) cross-species homologs. (**B**) Functional likelihood distribution of ITRs (left) and transcribed exons (right) with syntenic orthologs that are expressed (blue) or not (cyan). (**C**) Functional likelihood distributions of ITR (left) and transcribed exons (right) that are single copy (red) or duplicated (dark gray). (**D**) Functional likelihood distribution of ITRs (left) and transcribed exons (right) with duplicates that are expressed (blue) or not (cyan). (**E**) Boxplots of the relationship between functional likelihood and nucleotide substitution rate for ITRs (top left), transcribed pseudogenes (top right), transcribed exons (bottom left) and transcribed introns (bottom right).

Among duplicated ITRs, younger duplicates tend to have lower functional likelihood scores (*p*<0.02; **Fig. 6E**), but the correlation is exceedingly weak (ρ=0.05), similar to those of pseudogenes (ρ=0.11, *p*=0.15; **Fig. 6E**). By comparison, transcribed exons and introns show stronger positive correlations between timing of duplication and functional prediction scores (ρ=0.36 and 0.21, respectively; both *p*<9×10^-21^; **Fig. 6E**). To address the question if the time scale examined may be too short (*K*≤0.08), we assessed functional predictions of the 11 rice ITRs with retained duplicates from the >70 MYA ρ or σ WGD events. Five of these sequences (46%) were predicted as functional based on the stringent T1, while the other six of the 11 ρ/σ ITR duplicates failed to be predicted as non-functional based on T2. Together with cross-species sequence and expression conservation patterns, ancient duplicate retention provides strong support for the utility of our computational models for assessing sequence functionality and reaffirm our assertion that the majority of ITRs have pseudogene-like properties and likely represent noisy transcription. Meanwhile, a relatively small subset of ITRs with high functional likelihood may represent novel genes that warrant further study.

## DISCUSSION

In this study, we investigated the cross-species and post-duplication evolutionary characteristics of intergenic transcribed regions (ITRs) in four grass species. ITR sequences are primarily species-specific and, when ITR orthologs can be identified, the majority are not expressed. Similarly, ITR paralogs, which tend to be from recent duplication events, are usually not expressed. Thus, expression of an orthologous or duplicated ITR either evolved post-speciation/duplication or ITR orthologs/paralogs tend to quickly lose ancestral expression states. Either case suggests that expression among ITRs and related sequences may evolve more rapidly compared to the significantly higher degrees of cross-species and duplicate expression conservation seen among exons. We also found only rare examples of ITRs with paralogs retained from whole genome duplication events occurring ~12-70 MYA, suggesting there may be little selective pressure on most ITRs exerted through, for example, neofunctionalization. In addition to evolutionary characteristics, we evaluated the transcriptional features and found that ITRs tend to exhibit weak and narrow expression compared to annotated protein-coding exons. Overall, the evolutionary and expression characteristics explored here are consistent with the notion that intergenic expression is not subjected to strong selection and primarily represents transcriptional noise.

We also established robust functional prediction models for rice transcribed sequences using 44 evolutionary and biochemical characteristics of benchmark phenotype genes and expressed pseudogenes. These models distinguished between phenotype genes and pseudogene sequences with high accuracy. Based on model predictions, 583 ITRs in rice (8%) were classified as likely functional based on a conservative estimate of functionality (FPR of 5%). A previous study in *Arabidopsis thaliana* that employed selection within and between species as a criterion for defining likely functionality also found ~5% of the sequences functional (Moghe et al, 2013). In addition, in *A. thaliana*, a similar data integration framework and conservative functional estimate found that 7% of *A. thaliana* ITRs in were predicted as functional (Lloyd et al., 2018), suggesting that this proportion may represent a reasonable expectation for the proportion of potentially functional ITR sequences in plants.

Results from function prediction models indicated that ITRs with two characteristics: (1) cross-species expression conservation and (2) ancient retained duplicates primarily resemble phenotype genes. More importantly, prediction models also classified 203 (3%) of rice ITRs that lack sequence conservation as functional. This finding indicates that the data integration framework applied here can identify ITRs that may function in species-specific roles or when sequence conservation is not readily detectable (Ponting 2017). These two possibilities are not mutually exclusive. Ultimately, the majority of ITRs lack similarity to benchmark functional sequences. Considering that non-functionality cannot be tested but functionality can, we reiterate our argument that non-functionality should be treated as null hypothesis that is rejected when evidence suggests otherwise (Moghe et al 2013; Lloyd et al, 2018). Based on this logic, for most ITRs we cannot reject the null hypothesis that intergenic transcription primarily represents non-functional, transcriptional noise.

## METHODS

### Gene, pseudogene, and random intergenic annotation

The MAKER-P (r1065) genome annotation pipeline was used to reannotate the *Oryza sativa* (rice) Nipponbare (IGRSP-1.0 v7), *Brachypodium distachyon* (v1.2) and *Sorghum bicolor* (sorghum; v2.1) genome assemblies as previously described (Cantarel et al. 2008; Campbell et al. 2014). Repeats were masked using default parameters in RepeatMasker (v 4.0.3) for the sorghum and *B. distachyon* genomes, and a custom repeat library was created for rice using a method described previously (Campbell et al. 2014). RepeatMasker was run within the MAKER pipeline to mask repetitive elements. To aid in gene prediction, EST evidence was provided by transcriptome assemblies generated from selected publicly available data from the National Center for Biotechnology Information Sequence Read Archive (NCBI-SRA) (**Table S6**) using Trinity (version 2014) with a minimum contig length of 150 bp and the Jaccard clip option (Haas et al. 2013). Protein evidence was provided by the SwissProt plant protein dataset (ftp://ftp.uniprot.org/pub/databases/uniprot/current_release/knowledgebase/taxonomic_divisions/uniprot_sprot_plants.dat.gz), with the rice, sorghum, or *B. distachyon* protein sequences removed. For *Zea mays* (maize; v. 3.21), we utilized the MAKER-P gene annotation described by Law et al. (2015). Pseudogenes were identified in each species using a pipeline similar to that described by Zou et al. (Zou et al. 2009). We considered genome regions with at least 40 contiguous ambiguous nucleotides (Ns) as likely-unmappable, a length that represented the size of reads in RNA-sequencing (RNA-seq) datasets used in our analyses (see below). Likely-unmappable regions were excluded when calculating the size of total exon, intron, pseudogene, and intergenic space in **Fig. 1B**. Random sets of intergenic coordinates with equal length distributions as intergenic transcribed regions (ITRs, see below) were identified in each species. Random intergenic coordinates were sampled so that they did not overlap one another, transcribed regions, or likely-unmappable genome regions (as described above). We further filtered random intergenic coordinates to remove those that contained an ambiguous nucleotide, a step that removed 4-9% of random coordinates in each species.

### Identification and classification of transcribed regions

Multiple developmentally-matched RNA-seq datasets from rice, *B. distachyon*, sorghum, and maize (n = 11, 11, 10, and 14 respectively) were retrieved from the Sequence Read Archive at the National Center for Biotechnology Information (**Table S1**) (Davidson et al. 2011; Davidson et al. 2012). Reads were trimmed of low-scoring ends and Illumina adapter sequences using Trimmomatic v.0.33 (LEADING:3 TRAILING:3 SLIDINGWINDOW:4:20). Reads ≥20 nucleotides in length and with an average Phred score ≥20 were mapped to associated genomes using TopHat v.2.1.0 (-i 5000; -I 50000; all other parameters default) (Kim et al. 2013). Transcribed region coordinates were identified by assembling unique-mapped reads with Cufflinks v.2.2.1 (- min-intron-length 5000; -max-intron-length 50000; -m 150; --no-effective-length-correction) (Trapnell et al. 2010) while providing an associated genome sequence via the -b flag to correct for sequence-specific biases. Transcribed regions from across datasets that overlapped with one another by at least 1 nucleotide were merged. Merged transcribed regions were classified based on overlap with annotated exon, intron, and pseudogene sequences through a priority system: exon > intron > pseudogene (e.g. a transcribed region that overlapped both an exon and an intron was classified as exon). Transcribed regions that did not overlap with any gene or pseudogene annotation were classified as intergenic.

Transcribed regions were further categorized as likely protein coding or repetitive. Likely protein-coding transcribed regions were defined as those that contained a protein domain from Pfam v. 31.0 (Finn et al. 2016) in any translated frame or had significant translated sequence similarity to a plant protein annotated in Phytozome v.10 (BLAST v. 2.2.26; BLASTX E-value < 1×100^-5^) (Goodstein et al. 2012). Additionally, canonical RNA gene domains were identified within ITRs by searching for significant matches (E<1×100^-5^) with Rfam domain covariance models (v.12) (Nawrocki et al. 2015) using Infernal v.1.1 (Nawrocki and Eddy 2013)

A set of likely repetitive transcribed regions were defined based on the presence of repeat-associated Pfam protein domains or high numbers of duplicate sequences. To identify repeat-associated protein domains and set duplicate thresholds to call sequences as highly repetitive, a set of benchmark repetitive elements were identified in each unmasked genome using RepeatMasker v.4.0.5 (-nolow -norna -qq) with the default RepeatMasker repeat library and a custom library of rice repeats generated using previously described methods (Campbell et al. 2014). Transcribed regions that contained a Pfam v.31.0 protein domain that was significantly enriched among interspersed repeats relative to exon transcribed regions (Fisher’s exact test; adjusted *p*<0.05; Benjamini-Hochberg procedure) (Benjamini and Hochberg 1995) were considered likely-repetitive. We also identified a threshold based on the number of duplicate sequences required to call a sequence as highly-repetitive based on the duplicate counts of long terminal repeats (a subset of the interspersed repetitive elements identified by RepeatMasker) and transcribed regions that overlapped exons, through F-measure maximization. Resulting duplicate count thresholds to consider a transcribed region as repetitive were 10, 15, 33, and 202 for rice, *B. distachyon*, sorghum, and maize, respectively. For all duplication-related analysis, we excluded sequences that were classified as repetitive, as they were considered duplicated by definition.

### Sequence and expression conservation

Cross-species sequence matches were identified using BLAST searches (BLAST v. 2.2.26; BLASTN E-value <1×100^-5^) by searching nucleotide sequences of transcribed regions against repeat-masked whole genome sequences. The 95^th^ percentile E-values for random unexpressed sequences that lacked similarity to protein-coding or repetitive sequences were determined for all species: 1×100^-5^, 1×100^-6^, 1×100^-23^, and 1×100^-14^ for rice, *B*. distachyon, sorghum, and maize, respectively. A sequence was considered conserved if it had a cross-species match that was more significant than a 95^th^ percentile E-value. BLAST searches were performed within-species to identify duplicates and cross-species to identify putative homologs. Per base substitution rates (*K*) between within-species query and match sequences were calculated with PHAST (Hubisz et al. 2011) using a generalized time reversible model after aligning matched regions with EMBOSS-Needle v.6.5.7.0.

Expression conservation across species and following duplication was assessed by determining whether a cross-species or within-species match was overlapped by a transcribed region (as described above). Reciprocal matches between exon, intron, and intergenic transcribed regions for cross-species syntenic gene blocks (see below) were included in percent commonality calculations (**Fig. 3B,C**), which was calculated as the number of expressed tissues in common between a pair of transcribed regions divided by the total number of expressed tissues. The five tissues that were in common among all four species were considered: embryo, endosperm, seed, leaf, and anther. When replicate datasets were available, transcription evidence in a single dataset was required to consider a sequence expressed in the tissue. Similarly, expression in either seed datasets – 5 days after pollination (DAP) or 10 DAP – was required.

Expected percent commonality is affected by the expression breadth of two transcripts. For example, two orthologous transcribed regions that are both expressed in four out of five tissues have a minimum % commonality of 60%, while two orthologous transcribed regions both expressed in a single tissue have 20 possibilities for 0% commonality. As transcribed exons are expressed in more tissues than ITRs or transcribed introns (**Fig. S1C**), their % commonality cannot be directly compared. Instead, a subset of orthologous transcribed exons with expression breadth that matched ITR and transcribed intron pairs was selected to provide a direct comparison. For each orthologous ITR pair, pairs of orthologous transcribed exons that showed the same expression breadth as the ITR pair were identified. For example, in an orthologous ITR pair, if one sequence was expressed in one tissue and the second sequence in was expressed in two tissues, all orthologous transcribed exon pairs where one sequence was expressed in one tissue and the second was expressed in two tissues were identified. From this expression breadth-matched set of orthologous transcribed exon pairs, two pairs were randomly selected and used for % commonality calculations (**Fig. 3C**). The same framework was used to generate direct transcribed exon comparisons for orthologous transcribed intron pairs (**Fig. 3D**). Note that because the exon comparison was tailored specifically to the expression breadth distributions observed in pairs of orthologous ITRs or transcribed introns, the distribution of % commonality among transcribed exons is distinct in **Fig. 3C** and **Fig. 3D**. Random expectations of percent commonality (**Fig. 3C,D**, Rn) were calculated by randomly selecting tissues to match the expression breadth of intergenic and intron transcribed region pairs. For each orthologous ITR and transcribed intron pairs, 25 random pairs of tissues were selected that matched the observed expression breadth. Similar to the comparisons generated among orthologous transcribed exon pairs, the random pairs were generated specific for the expression observed in ITR or intron pairs, and so the distribution of random % commonality is distinct for the two sequence types. The probability that a tissue would be selected for a random set was proportional to how often it appeared among transcribed regions with expression conservation.

### Identification of syntenic gene blocks

We identified cross-species and within-species syntenic blocks by identifying of sets of collinear genes using MCScanX v.2 (Wang et al. 2012). Multiple minimum gene pair and maximum gap parameters were tested when generating collinear blocks (**Table S7**) and we utilized values of 10 and 10, respectively, for within-species collinear blocks and 10 and 2, respectively, for cross-species blocks. Cross-species blocks were identified between the more-closely-related species pairs: rice/*B. distachyon* and maize/sorghum. The rates of synonymous substitutions (*Ks*) between homologous protein-coding genes anchoring within-species syntenic blocks were calculated using the yn00 package in the Phylogenetic Analysis by Maximum Likelihood software (Yang 2007). Syntenic blocks with a median *Ks* ≥ 0.7 across all anchor genes in the syntenic block in all four species were considered associated with the ρ / σ whole genome duplication (WGD) events (Paterson et al. 2004; Tang et al. 2010), while syntenic blocks in maize with a median *Ks* <0.7 were associated with the more recent 12 MYO maize WGD (**Fig. S4**) (Swigoňová et al. 2004). An additional 254 gene-pair syntenic block in rice was identified with a median *Ks* of 0.13, suggesting recent duplication. However, due to the uncertain nature of the origin and timing of the duplication event (Wang et al. 2011), this block was excluded from further analysis. For exon and intron transcribed regions, syntenic duplicates and orthologs were identified as BLASTN matches (E-value<1×100^-5^) that overlapped a homologous anchor gene. For ITRs, and random intergenic sequences, syntenic duplicates and orthologs were BLASTN matches that were present within corresponding block regions circumscribed by homologous anchor genes (**see Fig. 4B)**.

### Rice functional prediction features

#### Transcription activity features

Twelve transcription-related features were generated for use in rice function prediction models (**Table S4**). The first 9 features were FPKM values (referred to as expression level; Level in **Table S4**) from each of the 9 tissues represented in the RNA-seq datasets described above. For replicate datasets (leaf and endosperm), expression level was taken as the average FPKM value if a region was expressed in both replicate datasets, and the single FPKM value otherwise. If a transcribed region was not expressed in an RNA-seq dataset, expression level was set to 0. Two additional features were represented by the maximum expression level among all RNA-seq datasets and the median FPKM among datasets where a transcribed region was expressed. The final feature was the expression breadth of a sequence, represented by the total number of tissues in which a sequence was expressed. For the breadth calculations, both seed datasets (5 and 10 DAP) and inflorescence datasets (early and emerging) were considered as a single tissue.

#### Sequence conservation features

Three sequence conservation-related features were generated. The first feature was the minimum BLASTN E-value to a sequence in *B. distachyon*, sorghum, or maize. A threshold of 1×100^-5^ was required to consider a match significant. Sequences without a significant cross-species match where given a score of 0. The second sequence-conservation feature was based on blocks of conserved nucleotides present across all four species were identified using the LASTZ/MULTIZ paradigm with rice as the target genome (Blanchette et al. 2004; Harris 2007; Hupalo and Kern 2013) (http://genomewiki.ucsc.edu/index.php/Whole_genome_alignment_howto). Among all conserved nucleotide blocks (CNBs; n=60,801), we identified those that were exonic (n=13,616), intronic (n=125), or intergenic (n=525) in all four species and were overlapped only by transcribed regions of the associated type (e.g. a conserved nucleotide block that was intergenic in all four species and was only overlapped by ITRs). The second conservation feature was the proportion of a sequence that overlapped a CNB. phastCons scores were calculated for each rice nucleotide within a conserved nucleotide block (Siepel et al. 2005), with conserved and non-conserved states estimated using the --estimate-trees option. The third sequence-conservation feature was the median per-base phastCons score across the proportion of a sequence that overlapped a conserved nucleotide block. If a sequence did not overlap a conserved block the phastCons score was set to 0.

#### Histone mark and nucleosome occupancy features

Twelve histone mark-related features among 10 histone marks were calculated based on 18 chromatin immunoprecipitation sequencing (ChIP-seq) datasets from NCBI-SRA (**Table S8**). Reads were trimmed as described above and mapped to the rice Michigan State University (MSU) v.7 genome with Bowtie v2.2.5 (default parameters). Spatial Clustering for Identification of ChIP-Enriched Regions v.1.1 (Xu et al. 2014) was used to identify significant ChIP-seq peaks with a non-overlapping window size of 200, a gap parameter of 600, and an effective genome size of 0.68 (Koehler et al. 2011). For datasets with control total histone or protein datasets, a false discovery rate (FDR) ≤0.05 was utilized (see **Table S8**). Peaks for histone marks with multiple datasets were merged. The first 10 histone mark-related features were represented by the percent overlap of a sequence with histone mark peaks for each histone marks (‘coverage’ in **Table S4**). The other two features were the number of activation-associated histone marks and repression-associated histone marks with a peak that overlapped a sequence. Eight marks were considered activation-associated and two were considered repression-associated (**Table S8**). A single nucleosome occupancy feature was also generated from micrococcal nuclease sequencing (MNase-seq) data. MNase-seq data was generated by Wu et al. (2014) and processed by Liu et al. (2015). The nucleosome occupancy feature was calculated as MNase-seq average read depth across the length of a sequence.

#### DNA methylation features

Sixteen DNA methylation features were calculated from bisulfite-sequencing (BS-seq) datasets from four tissues: embryo (SRR059000), endosperm (SRR059005), leaf (SRR618545), and panicle (SRR1520042). BS-seq reads were trimmed as described above and processed with Bismark v.3 (default parameters) to identify the number of reads that call cytosines as methylated and unmethylated in CG, CHG, and CHH (H = A, C, or T) contexts. The first 12 features were represented by the methylation level of a sequence in CG, CHG, and CHH contexts for each of the four tissues. Methylation level of a sequence was calculated as the number of reads that mapped to CG, CHG, or CHH sites that called a cytosine site as methylated divided by the total number of reads mapping to CG, CHG, or CHH sites within the sequence. A minimum of 5 reads across 5 cytosine sites were required to calculate methylation level. Multiple minimum read (range: 1-20) and site (range: 1-10) requirements were tested and found to have little effect on the ability of resulting methylation level features to distinguish between phenotype exons and transcribed pseudogene sequences (**Table S9**). The final four DNA methylation features were represented by whether a sequence exhibited a methylation pattern consistent with gene body methylation (GBM) in each of the four tissues. The GBM pattern is presence of CG methylation and absence of CHG and CHH methylation. Presence or absence of CG, CHG, and CHH states within a sequence were determined by binomial tests of the methylation level of a sequence (as described above) compared to the background methylation level across the whole genome for a given cytosine context. Binomial test *P*-values were corrected for multiple testing using the Benjamini and Hochberg method (FDR ≤ 0.05) (Benjamini and Hochberg 1995). Sequences that were significantly enriched in CG methylation relative to the genome background and not significantly enriched in CHG and CHH methylation were considered to be gene body methylated.

### Machine learning approach

The machine learning approach consisted of four steps: establishing a set of benchmark functional and nonfunctional sequences, identifying characteristics associated with benchmark sequences, integrating these characteristics via statistical learning methods to generate a functional prediction model, and applying function prediction models to all transcribed regions, including ITRs. We utilized the random forest algorithm implemented in the Scikit-learn software (Pedregosa et al. 2011) to perform machine learning runs aimed at distinguishing between functional and nonfunctional transcribed regions in rice. Benchmark functional transcribed regions were those that overlapped exons of genes with documented loss-of-function phenotypes (referred to as phenotype exons) (Lloyd et al. 2015; Oellrich et al. 2015). Benchmark nonfunctional sequences were represented by transcribed regions that overlapped pseudogene annotation (referred to as transcribed pseudogenes). We further filtered out pseudogenes that overlapped an exon from the MSU v.7 gene annotation. Benchmark non-coding sequences were transcribed regions that overlapped a set of high-confidence miRNA gene annotations from miRBase (referred to as transcribed miRNAs) (Kozomara and Griffiths-Jones 2014). Prior to model building, missing data points among features were imputed by sequential regression imputation implemented in the mice package in R (m = 1, seed = 500) (van Buuren and Groothuis-Oudshoorn 2011).

A machine learning-based prediction to distinguish between phenotype exons and transcribed pseudogenes was generated. We generated 100 balanced datasets that included equal proportions of phenotype exon and transcribed pseudogene sequences that were used for training and 10-fold cross-validation was implemented (i.e. 90% of a dataset was used for training and the held-out 10% used for testing). Parameter sweeps of maximum tree depth (3, 5, 10, and 50) and proportion of random features (10%, 25%, 50%, 75%, square root, and log2) values were performed, with 10 and 25% providing the highest performance, respectively. For each of the 100 balanced datasets, a prediction score for each transcribed region was generated that was equal to the proportion of 500 random forest trees that predicted the region as a phenotype exon; a final prediction score was calculated as the median of the 100 scores from the balanced datasets. Two thresholds to predict transcribed regions as likely-functional were generated based on a false positive rate of 5% (T1; score threshold = 0.60) and a false negative rate of 5% (T2; score threshold = 0.29). The Python script utilized to generate these predictions (ML_classification.py) is available on GitHub: https://github.com/ShiuLab/ML-Pipeline. Prediction models were also generated using the support vector machine and logistic regression algorithms implemented in Scikit-learn, but random forest provided the highest performance by AUC-ROC.

## ACKNOWLEDGEMENTS

The authors wish to thank Ning Jiang for providing a rice repeat library, Robin Buell for helpful discussions, and Jack R. S. DeCatters for inspiration and support. This work was partly supported by the National Science Foundation (NSF; IOS-1546617 and DEB-1655386) and the DOE Great Lakes Bioenergy Research Center (DOE Office of Science BER DE-SC0018409) to S.-H.S, NSF Graduate Research Fellowship to C.B.A., and the Michigan State University Dissertation Completion Fellowship to J.P.L.

## AUTHOR CONTRIBUTIONS

J.P.L., G.D.M., and S.-H.S. designed the research. J.P.L., M.J.B., K.L.C., and C.A.B. performed the research. J.P.L., G.D.M., M.J.B., K.L.C., C.A.B., and S.-H.S. wrote the article.

---

Fig. S1.Expression characteristics of transcribed regions in four Poaceae species. **(A)** Boxplots of length distributions among transcribed regions that overlap exon (Ex), intron (In), pseudogene (Ps), or intergenic (ITR) regions. Nt: nucleotides. Species abbreviation follows that of **Fig. 1**. **(B)** Boxplots of maximum FPKM distributions among all tissues. **(C)** Histograms of expression breadth (# of tissues with expression evidence) for transcribed regions. Ex (low): a subset of exons with expression levels ±5% of intergenic transcribed regions. **(D)** Percentage of transcribed regions that are reproducible across biological replicate leaf transcriptome datasets. Intergenic sequences were randomly-sampled to determine the background expected reproducibility (Rn).

Fig. S2. Ring plots displaying the number of species with evidence of expression for sequence blocks conserved across all four species. Sequence blocks shown were composed of only intergenic (**A**) or only exon (**B**) sequences from all species.

Fig. S3. Distributions of duplication types among transcribed exons (Ex, x-axis), ITRs, and random intergenic (Rn) sequences. Fisher’s exact tests were used to test significance between the proportion of sequences that were duplicated (sum of Ex, In, Ps, and ITR duplicate proportions) versus proportion of singletons. Although duplication rates between transcribed exons and ITRs could be significantly different, almost all differences were less than 5%, indicating that the presence of a duplicate is not informative to whether a sequence is likely part of an annotated gene. *: *p*<0.05, ***: *p*<0.001, n.s.: not significant, *p*≥0.05, In: intron, Ps: pseudogene.

Fig. S4. Distributions of synonymous substitution rates (*Ks*) between anchor genes of within-species collinear gene blocks. Circled numbers indicate *Ks* peaks associated with (1) a low-*Ks* large-scale duplication in rice, (2) a recent whole genome duplication (WGD) in maize, and (3) the ρ and σ WGD events. Average *Ks* values among ρ and σ duplicates have been estimated at 0.9 and 1.7, respectively (Paterson et al. 2004; Tang et al. 2010). *Ks* distributions for these two events are highly overlapping and cannot be effectively distinguished. Due to uncertain origin and timing of the low-*Ks* rice duplication (1) (Wang et al. 2011), duplicates from this event were not included in further analysis. *Os*: rice, *Bd*: *B. distachyon*, *Sb*: sorghum; *Zm*: maize.

Fig. S5. Boxplots of the relationship between distance to nearest gene and predicted functional likelihood for ITRs

Table S1.NCBI-SRA datasets used in transcribed region identification.

Table S2.Coordinates of intergenic transcribed regions in four Poaceae species.

Table S3.Machine learning instance coordinates and feature values.

Table S4.Machine learning feature list and single-feature AUC-ROC performance.

Table S5.Functional likelihood scores from machine learning predictions.

Table S6.NCBI-SRA datasets used in gene annotation.

Table S7.Parameters tested for syntenic block identification.

Table S8.NCBI-SRA datasets used in histone mark peak identification.

Table S9.Parameters tested for DNA methylation level feature calculation.

